# GeneHummus: A pipeline to define gene families and their expression in legumes and beyond

**DOI:** 10.1101/436659

**Authors:** Jose V. Die, Moamen Mahmoud Elmassry, Kimberly Hathaway LeBlanc, Olaitan I. Awe, Allissa Dillman, Ben Busby

## Abstract

During the last decade, plant biotechnological laboratories have sparked a monumental revolution with the rapid development of next sequencing technologies at affordable prices. Soon, these sequencing technologies and assembling of whole genomes will extend beyond the plant computational biologists and become commonplace within the plant biology disciplines. The current availability of large-scale genomic resources for non-traditional plant model systems (the so-called ‘orphan crops’) is enabling the construction of high-density integrated physical and genetic linkage maps with potential applications in plant breeding. The newly available fully sequenced plant genomes represent an incredible opportunity for comparative analyses that may reveal new aspects of genome biology and evolution. Analysis of the expansion and evolution of gene families across species is a common approach to infer biological functions. To date, the extent and role of gene families in plants has only been partially addressed and many gene families remain to be investigated. Manual identification of gene families is highly time-consuming and laborious, requiring an iterative process of manual and computational analysis to identify members of a given family, typically combining numerous BLAST searches and manually cleaning data. Due to the increasing abundance of genome sequences and the agronomical interest in plant gene families, the field needs a clear, automated annotation tool. Here, we present the GeneHummus pipeline, a step-by-step R-based pipeline for the identification, characterization and expression analysis of plant gene families. The impact of this pipeline comes from a reduction in hands-on annotation time combined with high specificity and sensitivity in extracting only proteins from the RefSeq database and providing the conserved domain architectures based on SPARCLE. As a case study we focused on the auxin receptor factors gene (ARF) family in *Cicer arietinum* (chickpea) and other legumes. We anticipate that our pipeline should be suitable for any plant gene family, and likely other gene families, vastly improving the speed and ease of genomic data processing.

## Introduction

Next-generation sequencing (NGS) has massively increased the number of nucleotide sequences deposited in public databases [1], revolutionizing plant sciences (Fig. 1). However, a bottle-neck in the field of plant sciences is the annotation of the proteins expressed from these sequences and the characterization of their functions. Such characterization can be used to improve agronomic performance and resistance/tolerance to specific environmental factors.

**Figure 1.**
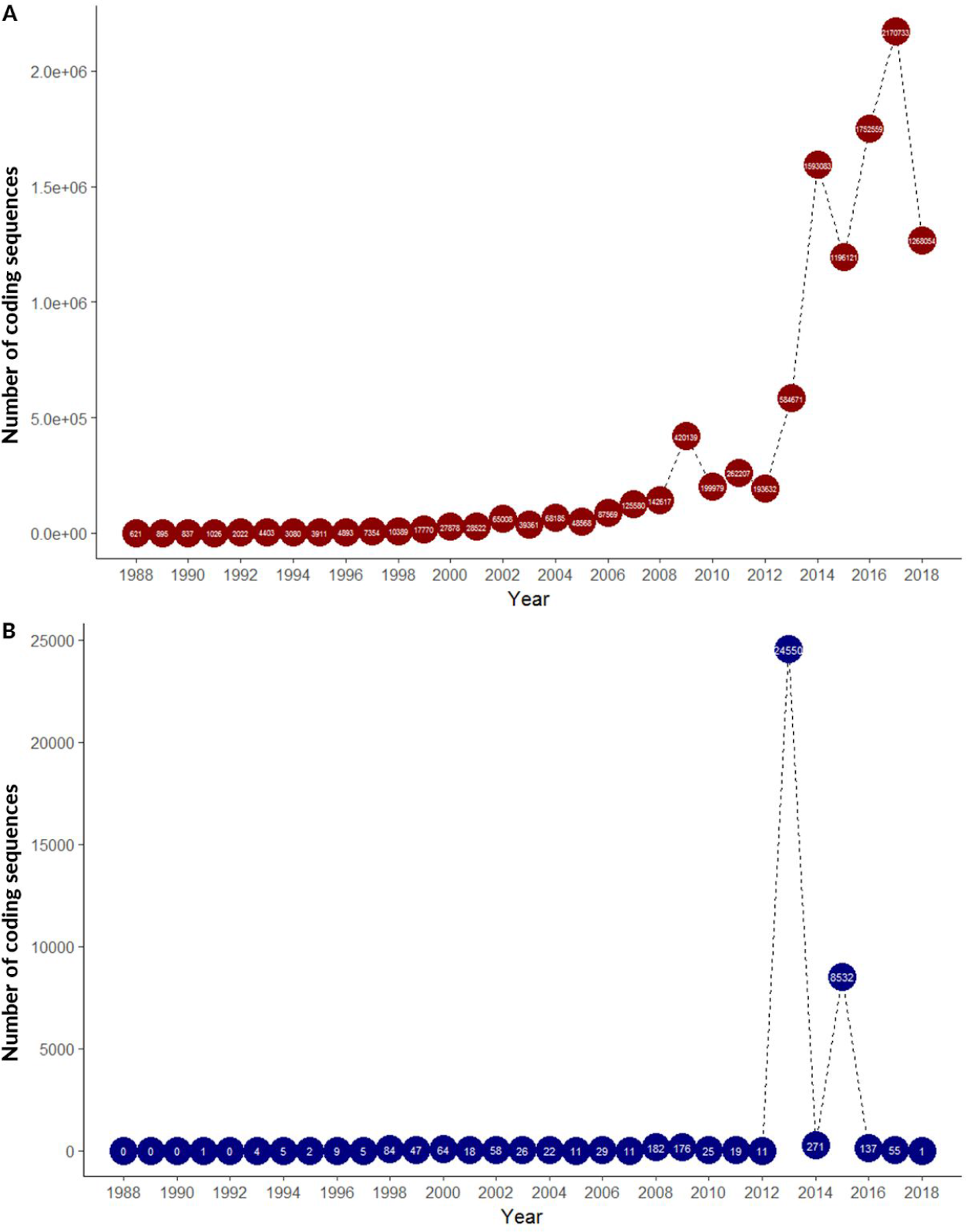
Statistics on the number of coding sequences deposited in NCBI. **(A)** Number of plant coding sequences deposited in NCBI over the past 3 decades. **(B)** Number of chickpea coding sequences deposited in NCBI over the past 3 decades.

The repetitive nature of identification and characterization of plant gene families is currently laborious and time-consuming. It requires an iterative process of computational analysis to identify the family members of a given family, based mainly on BLAST searches interspersed with manual curation and pruning. Due to the increasing number of sequences and the agronomical interest in plant gene families, this process clearly needs an automated tool.

The plant hormone auxin (indole-3-acetic acid) is a key regulator of virtually every aspect of plant growth and development [2]. As a central role of the auxin-signaling pathway, the auxin response factor (ARF) multigene family is present in all major divisions of land plants [3]. Considering the important role of ARF family members as regulators of plant growth and developmental processes, in the last few years there has been a considerable interest in studying the ARF family in both annual herbaceous plants [4, 5] and woody perennials [6, 7]. Characterization of ARF typically gives insights into the genomic structures [8], *loci* distribution across the genomes [9], sequence homology [10], phylogenetic history [11], and gene expression patterns [12] during development and/or biotic/abiotic stress.

Legumes comprise the third largest family of flowering plants and play an important agronomic role in the agriculture system of many countries through their contribution in the improvement of Nitrogen soil fertility by fixing atmospheric Nitrogen and minimising the use of inorganic nitrogen fertilizers [13]. These crops are grown in rotation with cereals contributing to a sustainable and environmentally friendly agriculture. Legumes are also important food crops for the food security with high protein content and other nutrients. In short, legumes including beans, chickpea, cowpea, lentils, pea, peanuts, and soybean are recognised by their highly nutritious source of protein and vital micronutrients, benefit people’s health and livelihoods, and enhance ecosystem resilience [14].

Here, we present the GeneHummus pipeline for the identification, characterization and expression analysis of plant gene families. As a case study we focused on the auxin receptor factors gene (ARF) family in *Cicer arietinum* (chickpea) and other legumes. For developing such a pipeline we used chickpea after having previously performed manual curation on the ARF gene family in this genome, so we had a gold standard dataset to which to compare our pipeline results. We anticipate that our pipeline should be suitable for any plant gene family, and likely other gene families, vastly improving the speed and ease of genomic data processing.

## Results & Discussion

### External validation of data

Numerous approaches have been developed to predict the function of different proteins. The Subfamily Protein Architecture Labeling Engine (SPARCLE; [15]), a recently developed resource by National Center for Biotechnology Information (NCBI), is one such approach. SPARCLE can help functional characterization of protein sequences by grouping them according to their characteristic domain architecture. We searched the SPARCLE database to obtain the whole set of molecular architectures based on the conserved domains that define the ARF gene family. Then, we filtered the data from SPARCLE to select the taxonomic group for the legume family (Fabaceae), the source database (RefSeq; [16]), and then by similarity score. After these filters, we obtained over 560 different ARF legume proteins encoded by ~330 gene *loci* (Fig. 2). After separating these results by species, our pipeline identified 24 ARF proteins in the chickpea genome (*Cicer arietinum*), reproducing the results obtained previously with an iterative exhaustive BLAST search [4]. These results validate the GeneHummus approach, which provides an automated way to produce these results in less than 6 minutes, as opposed to the exhaustive BLAST method which required significant manual curation and over 6 months of work. In addition, the GeneHummus pipeline also returned the number of ARF proteins in 9 other legume species in this 6 minutes processing window (Fig. 2). Interestingly, the number of ARF proteins is similar in different species within the same genus (*Arachis duranesis* and *Arachis ipaensis*; *Vigna angularis* and *Vigna radiata*), as may be expected. In addition, species known by possessing a high number of expanded paralogous genes by undergone whole-genome duplications events, such as *Glycine* and *Lupinus* lineages [17], showed the highest values both in number of transcripts and number of ARF *loci*. This increases confidence in the validity of our approach (Fig. 2). To find the ARFs in the different legume species using our pipeline, we developed an interactive shiny application to access this data https://genehummus.shinyapps.io/testshiny/. After running the GeneHummus pipeline, researchers interested in loading the relevant table for their gene families and taxa of interest can clone and modify https://github.com/NCBI-Hackathons/GeneHummus/blob/master/Shiny.R to easily share their results.

**Figure 2.**
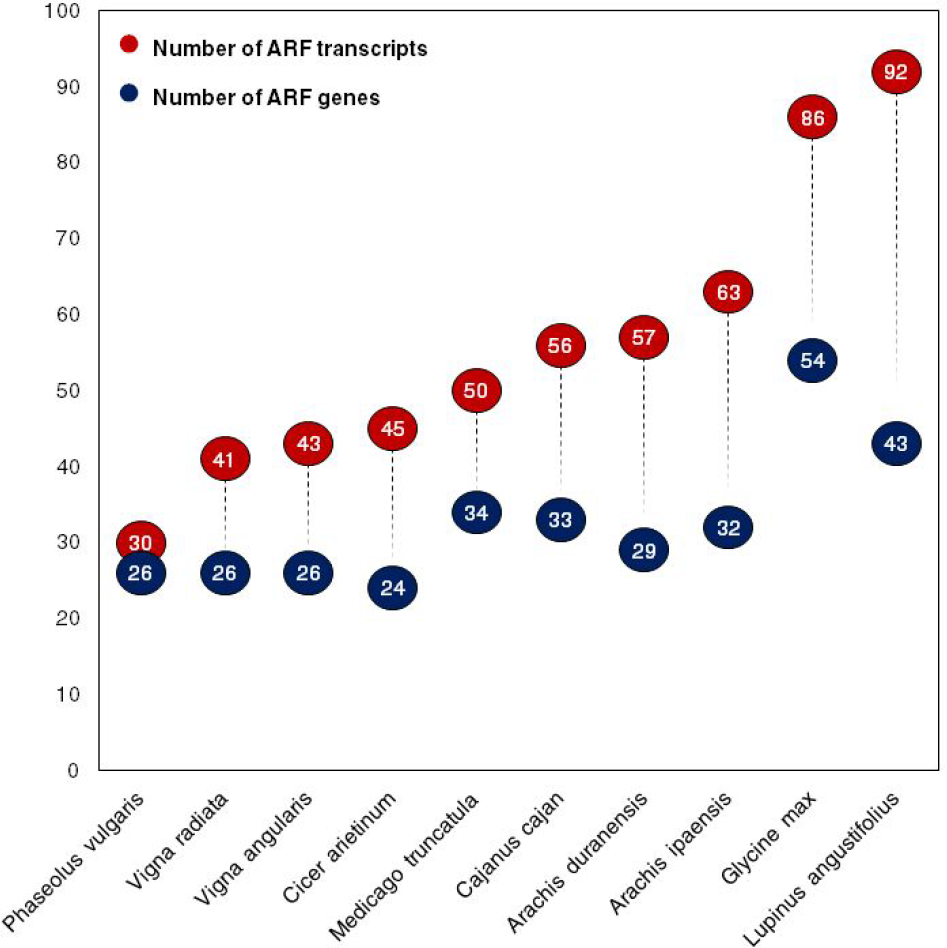
Total number of ARF sequences identified by CDART in legumes. Blue circles shows the number of ARF genes in each species of legumes, while red circles show the number of ARF transcripts.

### Phylogenetic Analysis

In addition to quantifying the number of ARF proteins in each legume species, the GeneHummus Pipeline also may be used for phylogenetic studies. Based on the conserved domains of ARF proteins, we explored and depicted the sequence relationships between the whole dataset. Two ARF proteins from the gymnosperm lineage were included as outgroup species. Gymnosperms have been resolved as the sister group of angiosperms. They diverged from their most recent common ancestor ~310 million years ago [18]. The phylogenetic distribution of the protein sequences revealed that all ARF sequences fall into two major groups (I and II) with well-supported bootstrap values (Fig. 3). The group I is the most numerous and may be further subdivided into clusters containing orthologs of the *Arabidopsis* sequences defining the well-known clades AtARF3/4-like, ARF12-like, ARF10/16-like, and ARF17-like [19]. Group II contain the cluster AtARF5-like. A second cluster in group II did not contain any *Arabidopsis* ortholog (data not shown) implying that this clade was derived through a long-term evolution for conserved functions across legume plant species. We labeled as sister pairs those proteins clustered together based on high bootstrap values (> 65%). Related to sister pairs involving chickpea, the phylogeny structures 8 sister pairs (seven pairs of *C. arietinum*-*M. truncatula* and one pair of *C. arietinum*-*L. angustifolius*). We did not observe any sister pair between two chickpea ARF proteins. This is an interesting evolutionary pattern. Chickpea diverged from *M. truncatula* ~ 10–20 million years ago [14]. Lack of chickpea sister pairs suggests that recent duplications (after chickpea and *Medicago* separated) have played a very limited role, if any, in the expansion of the ARF chickpea family, or that duplicated proteins did not change much since both species shared a common ancestor. Both hypothesis are plausible as well.

**Figure 3.**
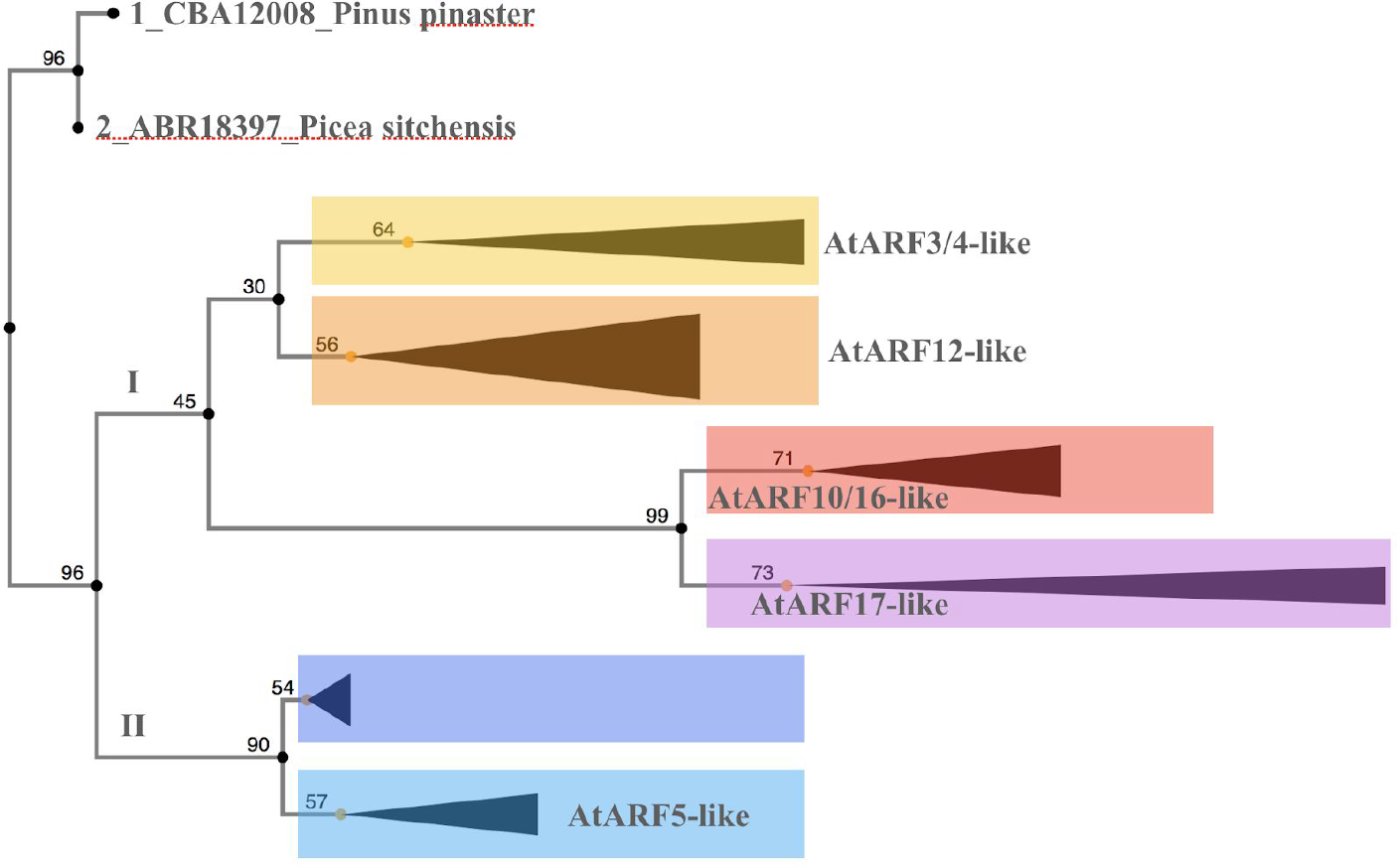
Analysis of phylogenetic relationships of ARF legume proteins. Phylogenetic analysis revealed two major clusters from the ARF lineage, one of which could be further subdivided into 6 clades The percentage of replicate trees in which the associated taxa clustered together in the bootstrap test (500 replicates) is shown next to the branches. Clades are named based on the phylogeny of the model plant *Arabidopsis thaliana*.

In addition, within the AtARF12-like group we observed a distinct clade made of 4 proteins based on bootstrap value, belonging to the ancestors of the cultivated peanut [20]. This suggests that these are orthologs of an ancestral gene emerged after the speciation of *Arachis* genus (Fig. 4). We are looking forward to the peanut genome becoming public so we can validate these results.

**Figure 4.**
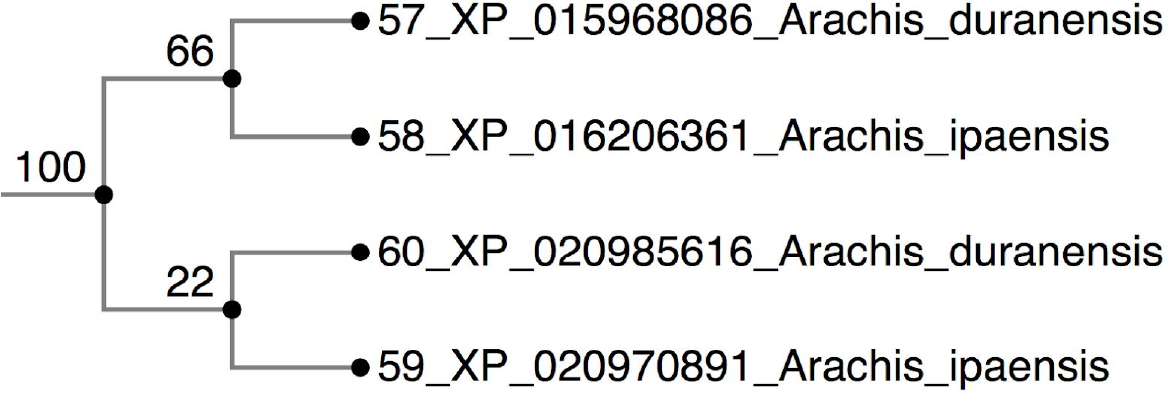
Cluster from clade AtARF12-like containing *Arachis* specific ARFs.

### Expression profiles: Gene Discovery

In addition to the identification of the ARF proteins by species and the phylogenetic analysis, GeneHummus can also incorporate data from SRA libraries using Magic-BLAST-based differential expression analysis to identify genes of interest for certain conditions. We identified datasets in the SRA that isolated sequences from the root of the chickpea plant raised in either drought or control conditions, and was identified as belonging to either a tolerant/drought or susceptible/drought strain. Upon analysis, we identified 3 transcripts that were differentially expressed in the drought-tolerant strain in drought conditions as opposed to the drought-susceptible strain (Fig. 5). This ARFs could be important targets for the genetic improvement of chickpea via conventional breeding or biotechnological approaches.

**Figure 5.**
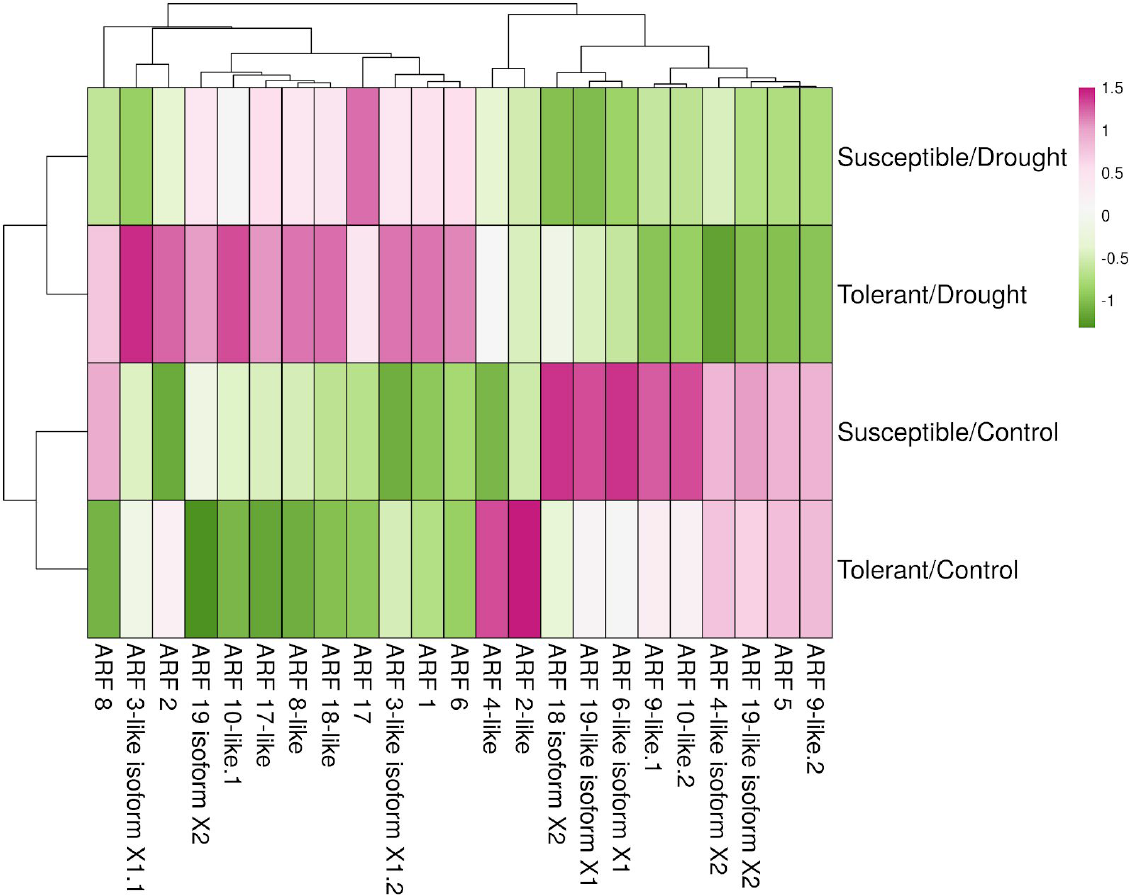
Differential abundance of ARF gene *loci in Cicer arietinum* under drought or normal conditions. Drought samples are clustered on the top,while control samples are clustered on the bottom.

## Conclusions

Applying the GeneHummus pipeline, we characterized the structure and phylogeny of the whole ARF proteins dataset in the legume family. As a case study, we also characterized the expression profile of the gene family in *Cicer arietinum*. The utility of this pipeline stems from a reduction in hands-on annotation time combined with high specificity and sensitivity in extracting proteins from the RefSeq database and providing interaction with the suite of other NCBI tools such as the conserved domain architectures based on the recently developed NCBI resources SPARCLE and the aligner Magic-BLAST. GeneHummus is a powerful tool for the identification of gene family sequences that could be used in phylogenetic analysis. Our results indicates that most proteins are very well conserved across genera, with abundant multi-species clades. This suggests that these proteins are involved in common basic cellular actions. This orthology information could be used to infer the function of a previously uncharacterized protein in a given species based on the known function of the protein in another genera. This is a particularly strong approach for comparative genomics. Once the sequences have been identified, given the ability of available SRA libraries for a number of tissues and conditions, the user can get the most out of the pipeline by using Magic-BLAST-based differential expression analysis to identify genes of interest for certain conditions. This tool will help investigators discover genes, and has particular applicability to plant breeding programs, among other applications. We anticipate that our pipeline should be suitable for any plant gene family and other gene families, vastly improving the speed and ease of genomic data processing.

## Methods

Documentation describing the ARF case study is available from our Github repository. (https://raw.githubusercontent.com/NCBI-Hackathons/GeneHummus/master/tutorial.md). This pipeline was designed for a biologist end-user with a minimal amount of programming experience, using open source and free software (R, NCBI tools) packaged inside a docker container -- a pre-built ready-to-run image containing all the software and the required configuration -- to guide the user through the identification, characterization and expression analysis of gene families. Plant gene families are characterized by common protein structure. The structure that defines a given family is known from literature. For example, the hidden Markov model (HMM) profiles of the ARF gene family are the B3 DNA binding domain (B3), AUX_RESP, and AUX/IAA, which correspond with the conserved domains Pfam 02362, Pfam 06507, Pfam 02309. The pipeline begins by defining the conserved domains accession numbers as a query (Fig. 6). Then, you can get the SPARCLE architectures for each conserved domain and extract only the labels that characterize the ARF gene family. Next, the protein ids for for each candidate ARF architecture are retrieved. Note that if you have a very long list of protein ids, you may receive a 414 error when you try to interact with the NCBI E-utilities. GeneHummus subsets the elements (up to 300 ids per list), so the functions can work properly. The retrieved protein ids are filtered by the taxonomy ids of interest [21], which in this case study were the Legumes ids. Genehummus returns only the protein ids hosted by the RefSeq database. At this point we have likely identified the whole set of ARF protein ids from the Legume family. Downloading the amino acid sequences from the target ids is straightforward. Once these sequences are downloaded, the relevant information may be used for phylogenetic and expression studies.

**Figure 6.**
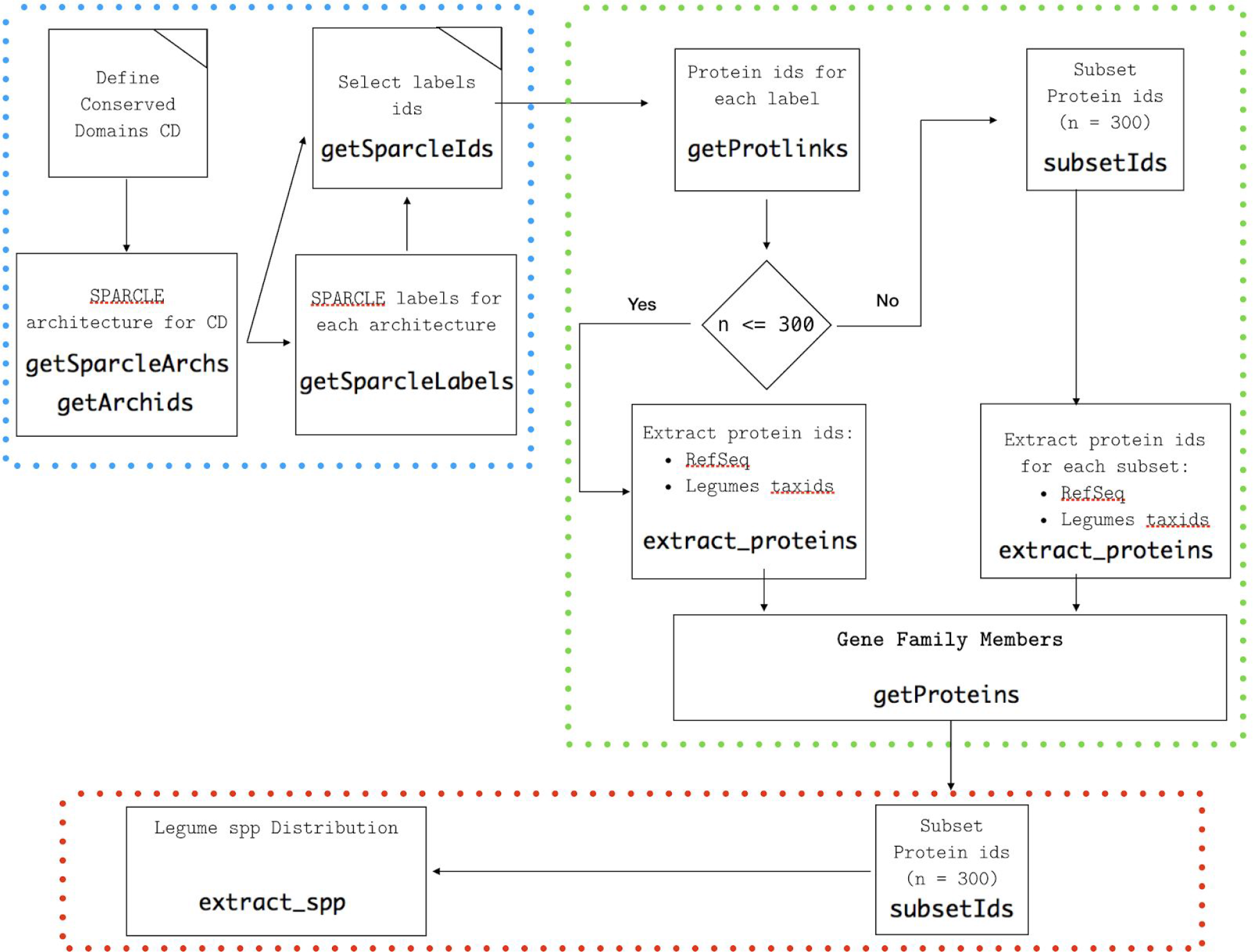
Workflow diagram for GeneHummus. The workflow shows the identification of protein families for legumes based on preparation of data (blue color), identification of family members (green color), and quantitative protein distribution per species (red color). R functions used in each step are highlighted in bold font.

### Phylogenetic Analysis

Multiple protein sequence alignments (MSAs) were performed on the conserved Pfam domains [Pfam 02309: AUX/IAA family; Pfam 06507: ARF (AUX_RESP); Pfam 02362: B3 DNA binding domain (B3)] for the whole dataset. Multiple alignments were performed with MAFFT version 7.402[22] [22] using standard methods (FFT-NS-i) and the following parameters: mafft --thread 10 --threadtb 5 -- threadit 0 --reorder --leavegappyregion. A recent online version of the software is available as well [23]. A NJ tree was conducted using the JTT substitution model, 500 replicates of bootstrap, and pair-wise detection of gaps. Two representative of gymnosperms (*Picea sitchensis* and *Pinus pinaster*) were included as outgroup species.

### Expression analysis

Using 1 single gene-model per locus, we created a BLAST database with the 24 ARF genes from the chickpea genome. Using Magic-BLAST [24], we studied the frequency of the ARF family in root tissues of two genotypes under drought stress and control conditions across 4 publicly available SRA libraries with the following parameters: alignment score = 125 bp, alignment identity ≥ 99% and read abundance in the four SRA libraries. A normalization factor was estimated for each SRA library by dividing the average SRA size by the corresponding SRA size. The normalization factor was applied to each read to give normalized counts.

### Implementation and Operation

Detailed instructions and dependencies can be found here: https://github.com/NCBI-Hackathons/GeneHummus/blob/master/tutorial.md

Briefly, R must be installed with the following libraries:

~~~
library(rentrez)
library(stringr)
library(dplyr)
library(curl)
library(httr)
~~~

To avoid errors with the HTTP2 framing layer, please run the following line:

~~~
httr::set_config(httr::config(http_version = 0))
~~~

The software can be run by importing the following functions:

~~~
source(“hummusFunctions.R”)
then loading the following file:
file = “data/ARFLegumes.RData”
load(file)
~~~

The software can then be extended by modifying the ARFLegumes.RData file to contain your taxids of interest.

The GeneHummus algorithm can also be run by running ./docker run ncbihackathons/genehummus3.

## Declarations

### Ethics approval and consent to participate

Not applicable

### Consent for publication

Not applicable

### Availability of data and materials

The datasets generated and/or analysed during the current study are available in the GeneHummus repository, https://github.com/NCBI-Hackathons/GeneHummus

## Authors’ contributions

JVD and MME participated in the design of the study, performed the bioinformatics and statistical analyses, generated figures and drafted the manuscript. KHL made substantial contributions to the bioinformatics analysis and the manuscript draft. OIA made contributions to the bioinformatics analysis and the manuscript draft. AD made contributions to the bioinformatics analysis. BB supervised the project and contributed to the manuscript draft.

All authors read and approved the final manuscript.

## Acknowledgements

We would like to thank Greg Boratyn, Aaron Marchler-Bauer and Lewis Geer from the NCBI MagicBlast and SPARCLE, and Jonathan Kans, the author of EDirect, for helpful discussions. We’d like to thank Steve Tsang for help with dockerizing R packages. We’d also like to thank Dan Smith, the lead administrator of the NCBI visiting bioinformatics program.

